# Local field potentials reflect cortical population dynamics in a region-specific and frequency-dependent manner

**DOI:** 10.1101/2021.05.31.446454

**Authors:** Cecilia Gallego-Carracedo, Matthew G. Perich, Raeed H. Chowdhury, Lee E. Miller, Juan A. Gallego

## Abstract

The spiking activity of populations of cortical neurons is well described by a small number of population-wide covariance patterns, the “latent dynamics”. These latent dynamics are largely driven by the same correlated synaptic currents across the circuit that determine the generation of local field potentials (LFP). Yet, the relationship between latent dynamics and LFPs remains largely unexplored. Here, we characterised this relationship for three different regions of primate sensorimotor cortex during reaching. The correlation between latent dynamics and LFPs was frequency-dependent and varied across regions. However, for any given region, this relationship remained stable across behaviour: in each of primary motor and premotor cortices, the LFP-latent dynamics correlation profile was remarkably similar between movement planning and execution. These robust associations between LFPs and neural population latent dynamics help bridge the wealth of studies reporting neural correlates of behaviour using either type of recordings.

## Introduction

Neuroscientists seek to understand how the brain drives behaviour. Researchers and clinicians often monitor brain activity using implanted electrodes^1–4^. Modern multielectrode arrays allow the spiking activity of hundreds of neurons to be observed simultaneously^5^. These same electrodes can also capture the lower frequency local field potentials (LFPs) that result from synaptic currents distributed across many thousands of neurons^6–10^. Uncovering a relationship between the collective dynamics of populations of single neurons and that of larger scale LFPs will help bridge studies looking at each of these signals in isolation, and advance our understanding of the human brain.

The spiking activity of neural populations can be well characterised based on the dynamics of relatively few covariance patterns^11,12^, which we refer to as the “latent dynamics”^13,14^. Studying these latent dynamics rather than the activity of single neurons has provided insight into how animals make decisions^15,16^, plan movements^17–19^, learn^20–24^, control movement timing^19,25,26^, and consistently perform the same behaviour^14^.

The latent dynamics are constrained by the neural covariance structure, which likely roughly reflects circuit connectivity^20,21,27^. Currents through these same connections are the main contributors to the generation of the LFPs^6–10^ (Figure 1A). In particular, correlations in the synaptic input currents across the population yield changes in LFP power at specific frequency bands, which often relate to specific brain functions^28–44^. For example, in the motor cortices, movement-related information lies primarily at low and high frequencies^36–39^ (i.e. <5 Hz and >50 Hz), yet, prior to movement initiation, there is a marked decrease in LFP activity in the less informative 15–30 Hz band^35,40,45,46^. In somatosensory cortical areas 2 and 3a, this same 15–30 Hz band contains signatures potentially related to afferent input from the limb^42^. It seems likely that these region-specific LFP phenomena relate to the neural population latent dynamics of each region, as they are both ultimately driven by the same synaptic currents.

**Figure 1:**
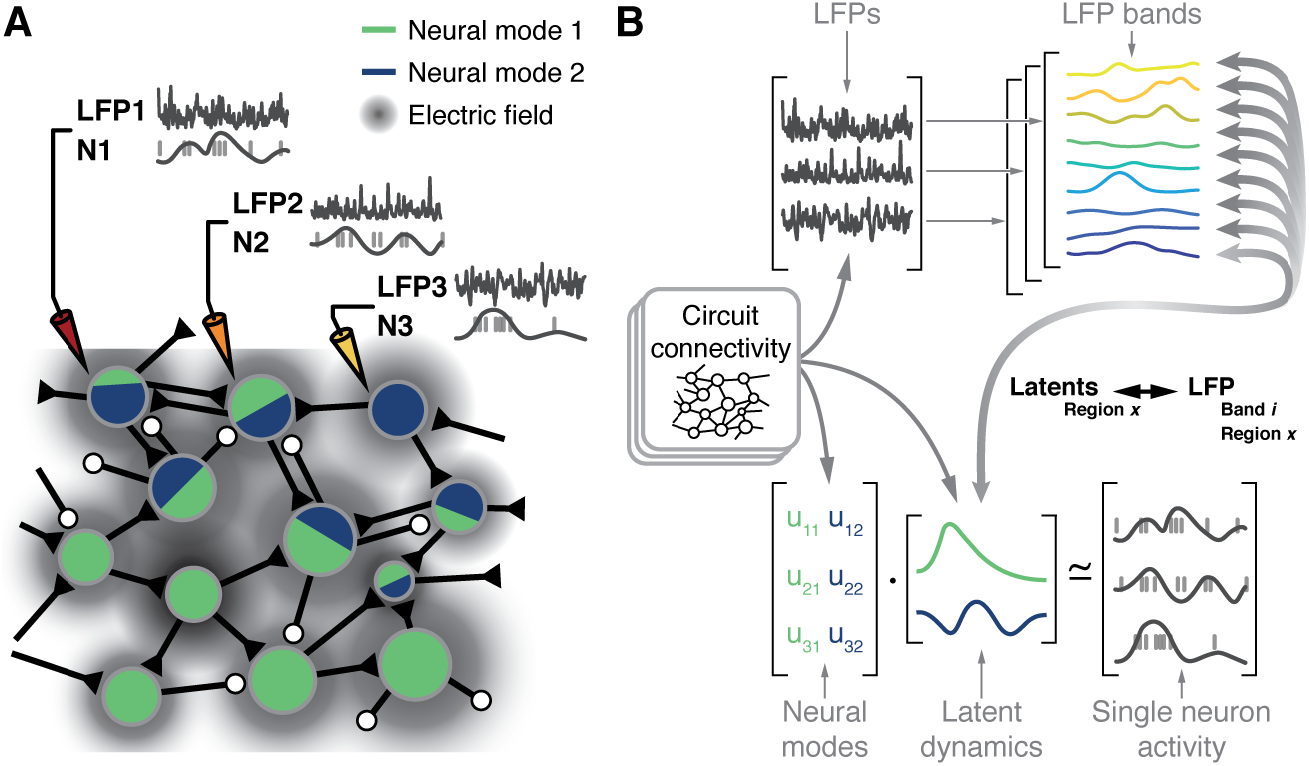
Hypothesis. **A.** Microelectrode arrays record both the spiking activity of single neurons (denoted as N1, N2, N3) and the local field potentials at these same sites (denoted as LFP1, LFP2, LFP3) of a cortical region X. **B.** Synchronization of synaptic currents within this circuit generate local field potentials in different bands (bottom). Circuit connectivity also constrains the coordinated latent dynamics of the N1, N2, N3 population (top). We thus hypothesize that, for each brain region, there will be a frequency-dependent association between LFPs and latent dynamics that should remain stable while the synaptic connectivity remains stable.

Previous studies investigated the relationship between single neuron activity and LFPs^31,40,43,47–50^. However, we lack a systematic description of how the different LFP bands relate to the latent dynamics reflecting the coordinated activity of the neural populations driving behaviour. Here, we report on the relationship between the different region-specific LFPs and the latent dynamics by addressing four hypotheses. First, since LFP^6–10^ and latent dynamics^20,21,27^ are shaped by the circuit biophysics (Figure 1B), we expect to find a robust relationship between these two signals. Second, this relationship should be frequency-dependent, because only specific LFP bands are strongly correlated with the behaviour^28–44^,^51^. Third, given that the synaptic connections remain stable in the short time scale spanning the preparation and execution of a movement, we anticipate the various LFP-latent dynamics associations would remain similarly stable throughout these two processes underlying behaviour. Lastly, since functionally different sensorimotor cortical areas have important differences in inputs^52–55^ and cytoarchitecture^56,57^, we hypothesize that the relationships between LFPs and latent dynamics would be region-specific.

We tested these four hypotheses using intracortical recordings from three different regions of the primate sensorimotor cortex during the same reaching behaviour: 1) primary motor cortex (M1), the main cortical output that controls movement execution^58–61^; 2) dorsal premotor cortex (PMd), a region that integrates inputs from many structures and is largely involved in movement planning^19,62–64^; and 3) area 2 of somatosensory cortex, where multimodal proprioceptive and cutaneous information about the state of the limb is processed^65–67^. Our results show that the relationship between the LFPs and the latent dynamics is largely frequency dependent and varies across cortical regions. Yet, the region-specific LFP-latent dynamics “correlation profiles” remain remarkably stable across the evolving processes mediating movement planning and execution^19,68,69^. Finally, we show that these relationships do not trivially arise from correlations between single unit firing rates and LFP recordings from the same intracortical electrode; instead, they reflect the coordinated activity of neural populations. This holds both during movement execution and as animals perform more “abstract computations” related to movement planning.

These results, which are critical for bridging studies of the sensorimotor system that use LFPs and neural population activity, paint a picture in which LFP bands relate to the latent dynamics in a stable, region-specific, and frequencydependent manner. This picture has one unexpected feature: some LFP bands are strongly correlated with latent dynamics yet carry little information about the animal’s movement. Given that low-dimensional latent dynamics have been found across many brain areas^12,70^, our findings are likely translated to beyond the sensorimotor system. Such translation may be especially insightful for areas largely studied based on LFP rhythms, such as the hippocampus^71^.

## Results

### Behavioural task and neural recordings

We trained four monkeys (C, M, H, and L) to perform an instructed delay centre-out reaching task using a manipulandum (Figure 2A) (Methods). The monkeys started each trial by holding a cursor in the central target before one of the eight outer targets was presented. After a variable delay period, an auditory “go cue” instructed them to move the cursor toward the presented target. A liquid reward was given after the monkeys had held the cursor in the target for a given period of time. For the monkeys with implants in area 2 (monkeys H and L), both the delay and holding periods were omitted.

**Figure 2:**
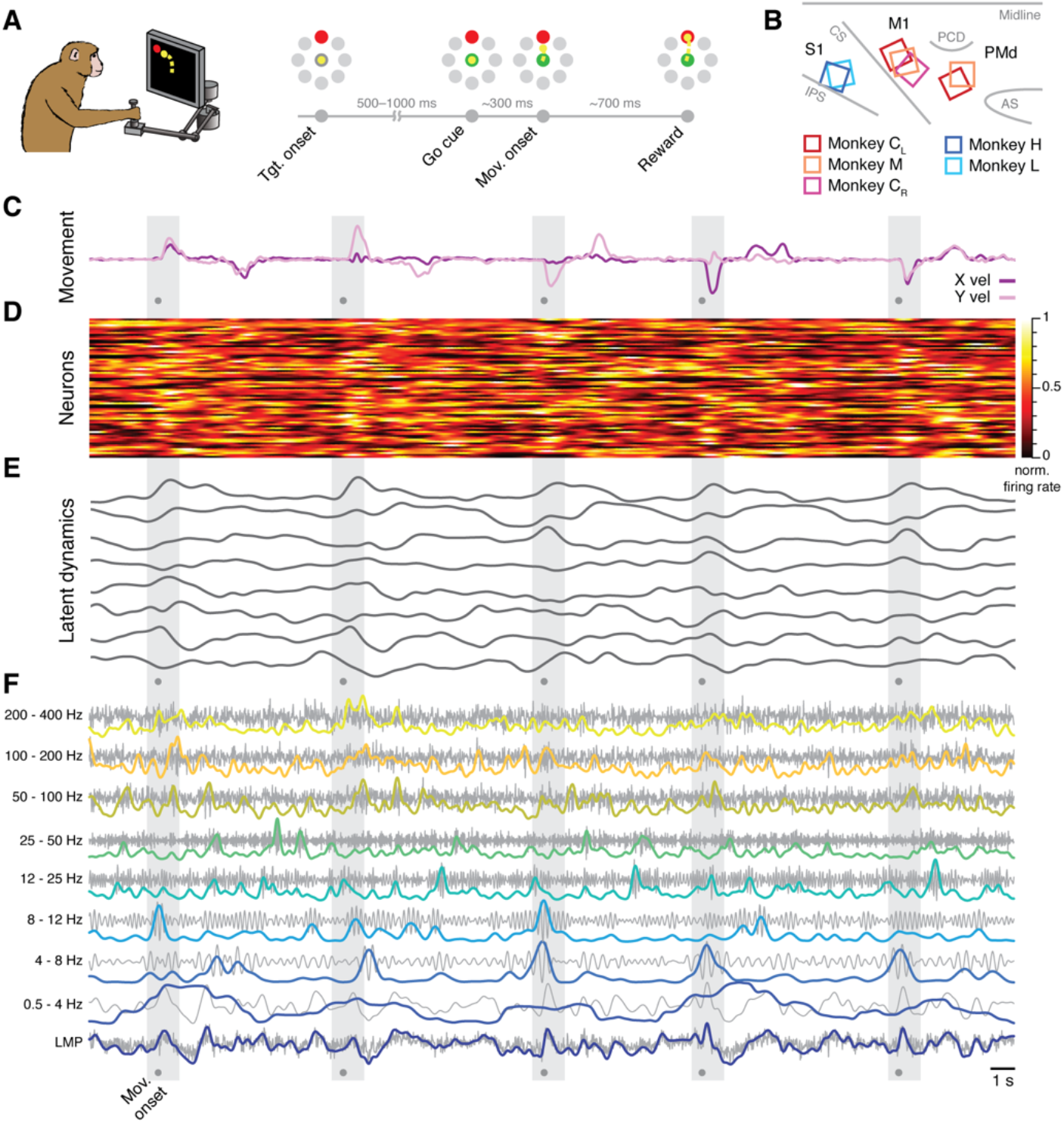
Behavioural task and neural recordings. **A.** Monkeys performed a reaching task using a planar manipulandum. **B.** Approximate locations of all seven arrays; each colour is one set of implants (legend). IPS, intraparietal sulcus; CS, central sulcus; AS, arcuate sulcus; PCD, precentral dimple. **C.** Hand velocity along the horizontal and vertical axes during five trials from a representative session from Monkey C_L_. Shaded grey areas, reach epoch for each trial; dots, movement onset. **D.** Example neural recordings showing the activity of 84 simultaneously recorded putative neurons. **E.** Latent dynamics corresponding to the neural firing rates in D. **F.** Example LFP recordings. Each row shows the LFP activity in one of the nine bands we studied (grey) along with its power (colour). Panels A, B adapted from Ref. 14.

Each monkey was implanted with one or two 96-channel microelectrode arrays (Figure 2B). Monkeys C and M had dual implants in the arm regions of M1 and PMd, whereas Monkeys H and L had a single implant in the arm region of area 2. We also analysed a second dataset from the other (right) hemisphere of monkey C, which we recorded in a different set of experiments; we denote this dataset as Monkey C_R_.

We simultaneously recorded neural spiking and LFP on each electrode. The spiking signals were manually sorted to identify putative single neurons (Figure 2C). To compute the neural population latent dynamics, we constructed a highdimensional neural state space in which the smoothed firing rate of each neuron was represented on a different axis; this way, the state of a population of *N* neurons at a given time *t* corresponds to one point in an *N*-dimensional state space. We used Principal Component Analysis (PCA) to find the lower-dimensional neural manifold spanning the dominant population-wide activity patterns^12,72^ choosing the following manifold dimensionalities based on previous studies^14,17,22^: 10 dimensions for M1, 15 for PMd, and 8 dimensions for area 2. We computed the latent dynamics by projecting the neural activity into each of the axes (PCs) that defined a given neural manifold (Figure 2D).

We calculated the LFP power in eight standard frequency bands^37–39,73^: 0.5-4 Hz, 4-8 Hz, 8-12 Hz, 12-25 Hz, 25-50 Hz, 50-100Hz, 100-200 Hz, 200-400 Hz for each electrode from which we could record at least one putative neuron. We also computed the local motor potential (LMP), which captures fluctuations in total LFP power^38,39^ (Figure 2E). For full details, see Methods section.

### The LFP-latent dynamics correlation profiles in primary motor cortex are frequencydependent

We first investigated the similarity between the latent dynamics in M1 and each of the LFP frequency bands defined above. We used Canonical Correlation Analysis (CCA), a method that quantifies the similarity between two sets of signals by finding the linear transformations that maximise their correlation^14,74,75^ (Methods); we refer to this process simply as “alignment.” For each session, we aligned the latent dynamics with the LFP power in each frequency band (Figure 3A). We performed this process separately for each recorded LFP signal, which yielded nine distributions of (canonical) correlation coefficients, one per frequency band, with as many samples as electrodes containing identified neurons (shown as individual data points in Figure 3A).

**Figure 3:**
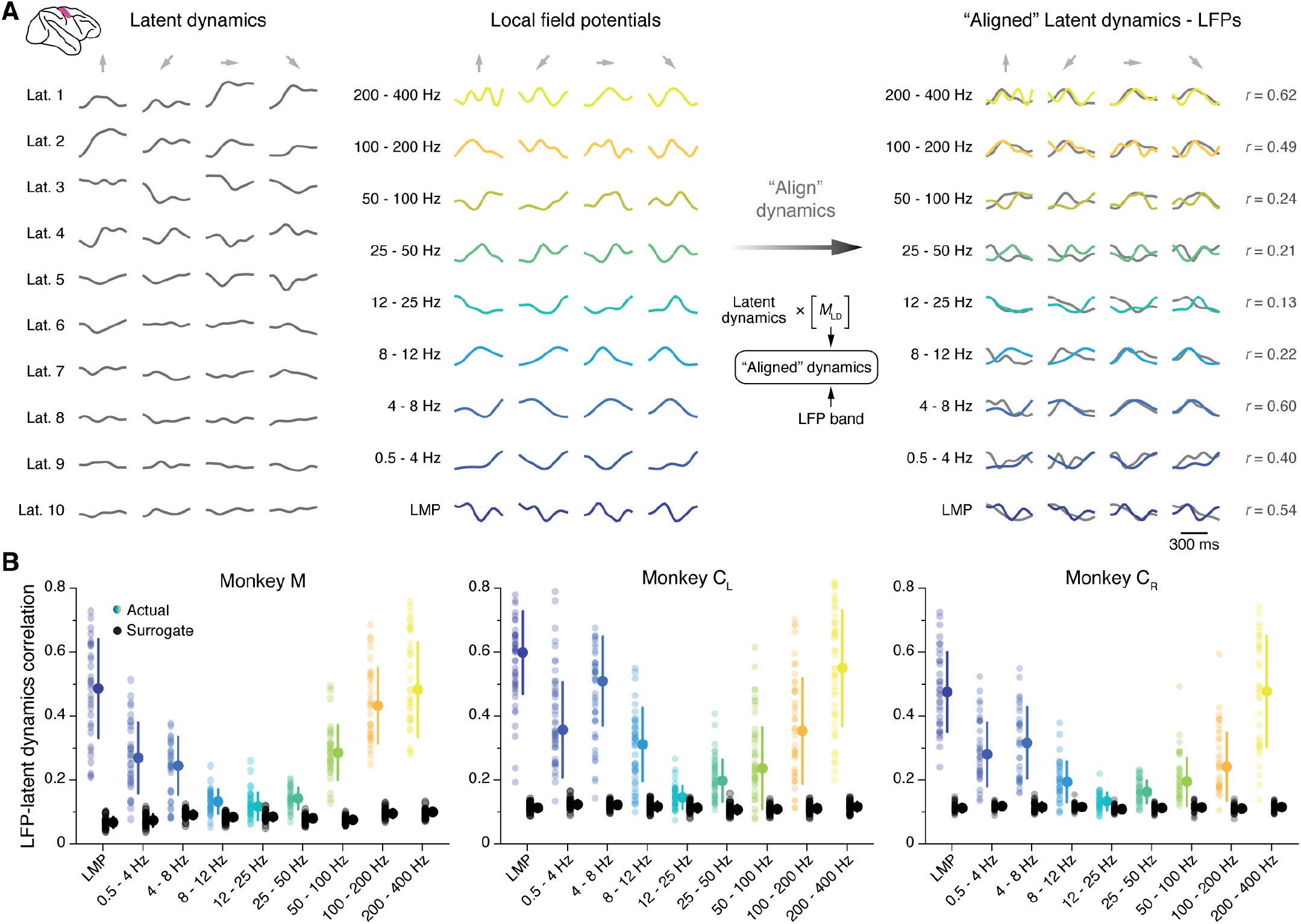
Similarity between M1 latent dynamics and each LFP band during movement execution. **A.** Left: Example M1 latent dynamics during three reaches to different targets (direction indicated by the arrows above each column); figure shows top ten dimensions of the latent dynamics. Middle: Example LFP power in each of the nine bands we study during the same three trials. Right: CCA “alignment” finds strong similarities between certain LFP bands and the latent dynamics. *r*, correlation coefficient. From one session from Monkey C_L_. **B.** Correlation between each LFP band and the latent dynamics (coloured markers) during one representative session from each M1 monkey. Black markers show the control correlation values obtained after generating surrogate neural activity using TME; note that at low and high LFP frequencies, the actual correlations are much larger than the surrogate correlations. Error bars, median ± s.d.

Figure 3B shows one representative session for each M1 monkey, indicating that the relationship between LFPs and latent dynamics was both consistent across subjects and frequency dependent: there were clear correlations between LFP and latent dynamics in the low and high bands (median correlation: 0.3–0.6), whereas the correlations approached zero in the mid-range bands (8–50 Hz) (coloured distributions in Figure 3B; Figure S1A summarizes all datasets).

We devised a control analysis to verify that the larger correlations of the low and high frequency bands captured a significant relationship between LFPs and latent dynamics. Using tensor maximum entropy^76^ (TME), we generated surrogate neural firing rates that both lay on the same neural manifold –i.e. preserved the covariance across neurons–, and had temporal statistics similar to the actual data (examples in Figure S1B,C) (Methods). We reasoned that despite their spectral similarity, the correlations between the LFPs and the surrogate latent dynamics should be much lower than those between the LFPs and the actual latent dynamics. As predicted, the surrogate correlations were significantly lower than the actual correlations (median ≤ 0.1; *P* <0.001, Wilcoxon rank sum test; compare the black and coloured distributions in Figure 3B). Thus, the difference in LFP-latent dynamics correlations across LFP frequencies could not be trivially explained by similar spectral characteristics between the signals. We obtained similar LFP-latent dynamics correlation profiles for different manifold dimensionalities (Figure S1D), and when pooling all putative single neurons on an electrode into “multi-units” (Figure S2A). Our results also held when modifying the specific frequency bands used for LFP preprocessing within reasonable intervals (Figure S2B), and could not be explained by differences in LFP variance across bands (Figure S2C).

M1 latent dynamics^13,14^ and some LFP bands^36–39^ allow accurate prediction or “decoding” of movement kinematics. We thus asked whether the strength of the LFP-latent dynamics correlation anticipates how well an LFP band predicts behaviour. We used standard linear decoders to predict hand velocity from each LFP band and the latent dynamics^14,38^ (Figure 4A) (Methods). Decoding accuracy varied considerably across LFP bands, with some (e.g., the LMP and 200– 400 Hz) being nearly as predictive as the latent dynamics (Figure 4B,C; Figure S3). Interestingly, the accuracy of each LFP band was strongly correlated with its similarity to the latent dynamics (R^2^ = 0.78; *P* < 10^-3^; see Fig 4D) even though all bands had similar variance (Figure S2C). Thus, for M1, the relationship between LFP power and latent dynamics varies across frequency bands and predicts how strongly each band relates to the ongoing motor output.

**Figure 4:**
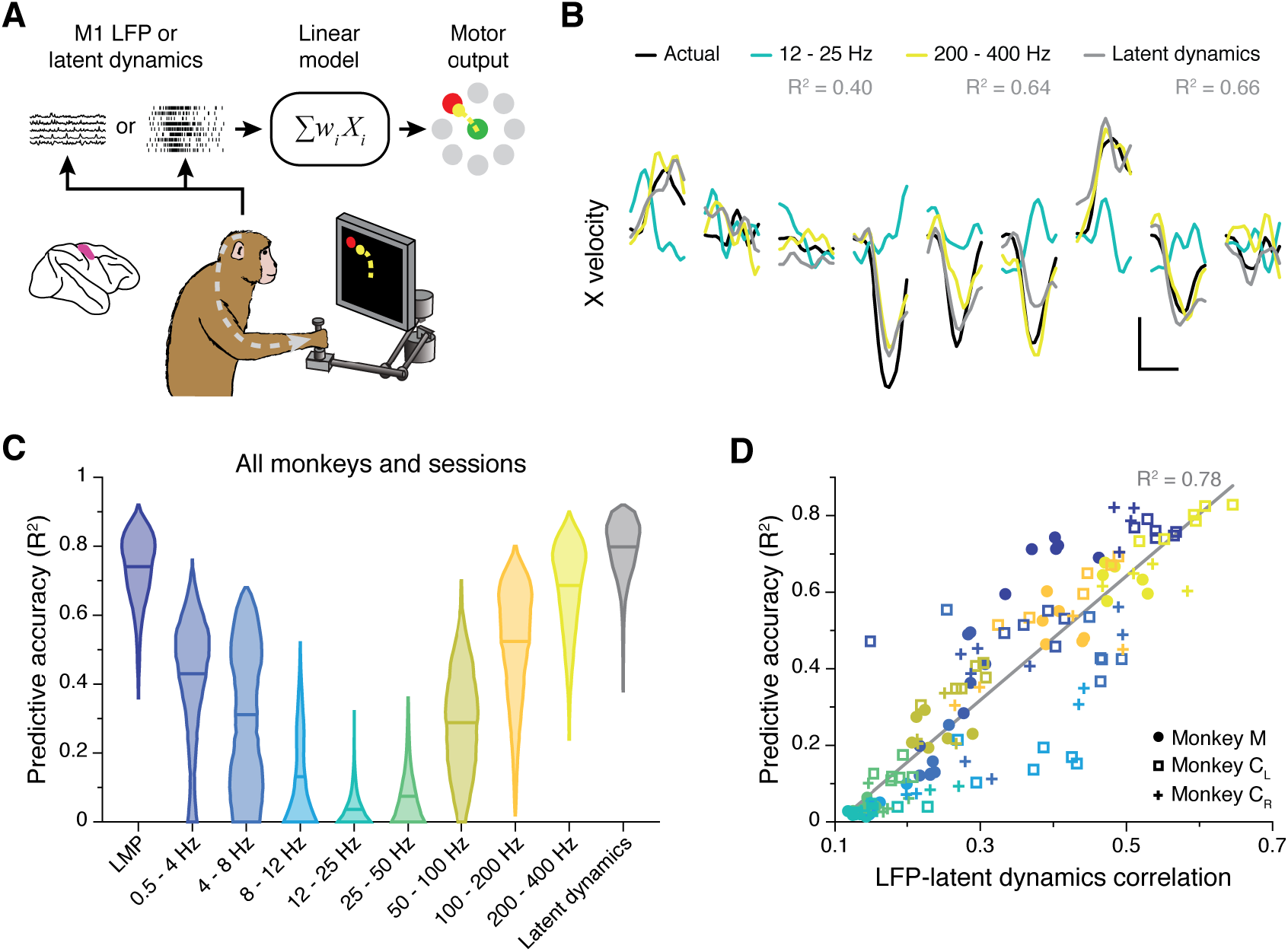
Decoding movement kinematics from M1 LFPs. **A.** We trained linear decoders to predict hand velocity in the x and y axes from the either the latent dynamics or the LFP power in different frequency bands. **B.** Example x-axis velocity predictions during nine randomly selected trials from one session from Monkey CL. Note the clear difference in predictive accuracy between the example LFP bands (values for each input signal during the example trials shown are indicated in the legend). Scale bars: horizontal, 300 ms; vertical, 10 cm·s^-1^. **C.** Predictive accuracy of each LFP band (colours) and the latent dynamics (grey) pooled across all M1 monkeys and sessions. Violin, probability density for one frequency band; horizontal bars, median. **D.** Linear relationship between the M1 LFP-latent dynamics correlation coefficients and LFP predictive accuracy. Each marker denotes one LFP band from one session; different monkeys are shown using different markers (legend). Markers are colour-coded as in C.

### The LFP-latent dynamics correlation profiles are stable between movement planning and execution

We identified a robust frequency-dependent relationship between M1 LFPs and latent dynamics during movement execution. Since M1 is also involved in movement planning^19,62,77^, we asked whether this relationship remains stable between these two processes underlying behaviour. Repeating the previous CCA alignment procedure for the instructed delay epoch showed that the LFP-latent dynamics correlation profile was virtually identical between movement planning and execution (Figure 5A): a correlation analysis between all pairs of LFP-latent dynamics correlations across all frequency bands, sessions, and monkeys revealed a strong linear association (R^2^ = 0.87; *P*< 0.001; Figure 5B).

To provide a reference for the observed correlations, we repeated the alignment procedure during the inter-trial interval, when monkeys were not actively planning or executing the behaviour. In this period, M1 neurons typically fire less (Figure S4A,B) and become less correlated (Figure S4C), likely due to the lack of strong synaptic currents driving the population. Since synaptic currents are the main contributors to LFP generation^9^, we predicted that the LFP-latent dynamics correlations should decrease during the inter-trial intervals. As expected, the correlations did indeed become much lower during the inter-trial intervals than during movement execution (Figure 5A,C).

**Figure 5:**
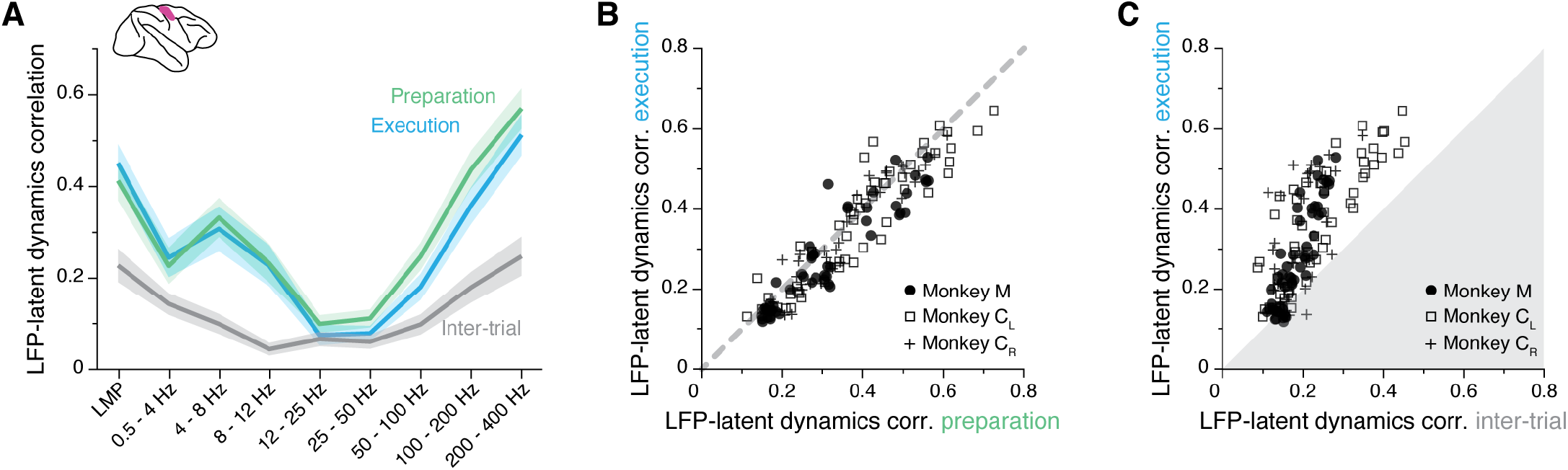
The M1 LFP-latent dynamics correlation profile is preserved between movement planning and execution. **A.** LFP-latent dynamics correlation profiles across all sessions from all three M1 monkeys. Line and shaded areas, mean ± s.e.m. Each epoch is shown in a different colour. **B.** Comparison between the LFP-latent dynamics correlations during movement preparation and execution. Each marker shows one frequency band for one session; each monkey is represented using a different marker (legend). Note the very strong similarity between epochs. **C.** Comparison between the LFP-latent dynamics correlations during movement execution and the inter-trial period. Data formatted as in B. Note the marked decrease in correlations during the inter-trial period.

The observation that the LFP-latent dynamics correlation profile remains preserved between movement planning and execution extends our previous findings to a more abstract process occurring in the absence of movement. It also indicates that the observed association is not a trivial epiphenomenal consequence of the frequency content of movement, but instead likely reflects underlying physiological processes related to the production of behaviour. The decrease in LFP-latent dynamics correlations during the inter-trial intervals provides additional support for this interpretation.

### The LFP-latent dynamics correlation profiles change between primary motor and premotor cortices

The previous sections demonstrated a frequency-dependent relationship between the LFPs and latent dynamics in M1. We next asked whether similar associations could be found in other sensorimotor cortical regions. We studied PMd, a “higher” motor region with a different cytoarchitecture from M1^57^ thought to be key for planning an action^19,64,68,78,79^. We repeated the comparison between latent dynamics and LFP bands first focusing on the movement planning epoch (Figure 6A). As for M1, the LFP-latent dynamics correlations were markedly frequency-dependent (Figure 6B), much stronger in the low and high frequencies, for which they greatly exceeded the surrogate control (Wilcoxon rank sum, *P*<0.001). Interestingly, virtually all the datasets (10 of 12; pooled results in Figure S5A) exhibited a strong correlation at 12–25 Hz, which was notably absent in M1.

Similar to M1, PMd is involved in both movement preparation and execution. Thus, we tested whether the LFP-latent dynamics correlation profile was preserved as in M1, and found it to be stable between movement planning and execution (R^2^ = 0.79; *P*< 0.001; Figure 6C). During the inter-trial period, when PMd neurons become less active, the correlations dropped, though less markedly than for M1 (Figure 6D). Therefore, the frequency dependent relationship between LFPs and latent dynamics changes across cortical areas but, within an area, remains stable across different behaviour-related processes those areas are involved in.

**Figure 6:**
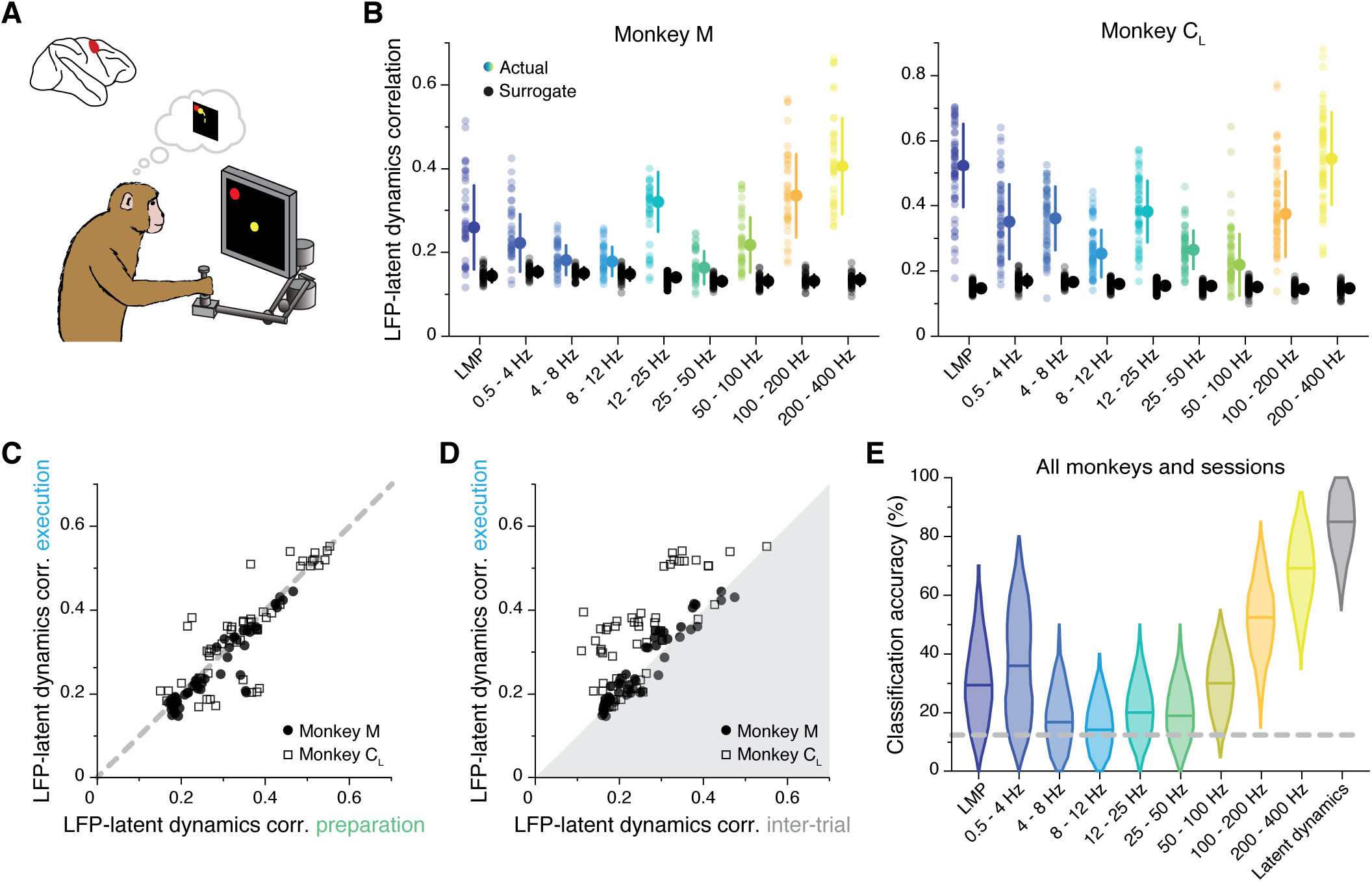
Similarity between PMd latent dynamics and each LFP band during movement preparation. **A.** We focus on the instructed delay period following target presentation and preceding the subsequent go cue**. B.** Correlation between each LFP band and the latent dynamics (coloured markers) during one representative session from each PMd monkey. Black markers show the control correlation values obtained after generating surrogate neural activity using TME. Error bars, median ± s.d. **C.** Comparison between the LFP-latent dynamics correlations during movement preparation and execution. Each marker shows one frequency band for one session; each monkey is represented using a different marker (legend). Note again the very strong similarity between epochs. **D.** Comparison between the LFP-latent dynamics correlations during movement execution and the inter-trial period. Data formatted as in C. **E.** Accuracy of classifiers that predict the reach direction based on each LFP band (colours) and the latent dynamics (grey); data pooled across all PMd monkeys and sessions. Violin, probability density for one frequency band; horizontal bars, median; dashed horizontal line, chance level.

Last, we studied whether the LFP’s ability to predict the upcoming movement^80^ was also frequency dependent. The performance of classifiers that predicted the upcoming target from the instructed delay activity^14,63^ (Methods), varied dramatically across LFP bands (Figure 6E; individual examples in Figure S5B): high-frequency bands were almost as accurate as the latent dynamics, while classifier accuracy at low-frequencies was worse and quite variable across monkeys (~50% accuracy for Monkey C; close to chance for Monkey M). Interestingly, not all bands that were significantly correlated with the latent dynamics also predicted the upcoming target (Figure S5C); most notably, despite being among the most correlated bands, 12–25 Hz prediction accuracy was barely above chance.

### The LFP-latent dynamics correlation profiles are different for area 2 of primary somatosensory cortex

While LFPs in PMd and M1 primarily relate to preparing and generating movement, oscillations in area 2 of somatosensory cortex likely reflect somatosensory feedback processing^46,81^, perhaps in combination with efference copy signals^42^. We thus asked whether the LFP-latent dynamics correlation profile in area 2 is different from that of motor regions (Figure 7A). Both the low and high frequency LFP bands of area 2 were strongly correlated with the latent dynamics (Figure 7B; Figure S6A), similar to the profile shown above in both M1 and PMd. Intriguingly, we observed a strong, significant correlation at 12–25 Hz (and for Monkey H also at 25-50 Hz), which we previously observed in PMd but not in M1. Again, all LFP-latent dynamics correlations dropped dramatically during the inter-trial period (Figure 7C), when monkeys did not move, and area 2 neurons go practically silent^66^.

**Figure 7:**
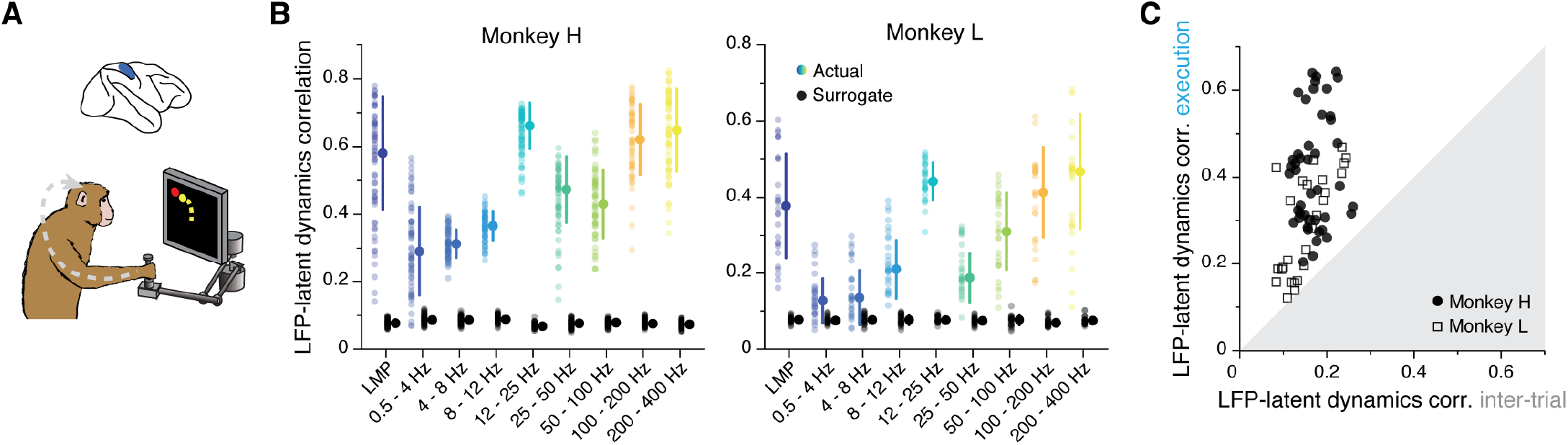
Similarity between area 2 latent dynamics and each LFP band during feedback processing. **A.** We focused on movement execution, when area 2 receives proprioceptive input about the state of the limb. **B.** Correlation between each LFP band and the latent dynamics (coloured markers) during one representative session from each area 2 monkey; same format as Figures 3B and 6B. Note the clear frequency-dependent relationship between LFP and latent dynamics. Error bars, median ± s.d. **C.** Comparison between the LFP-latent dynamics correlations during movement execution and the inter-trial period; same format as Figures 5C and 6D. Note the marked decrease in LFP-latent dynamics correlation during the inter-trial period.

As in the other regions, we examined whether the ability to make predictions of the monkey’s behaviour from each LFP band was correlated with the strength of its association with the latent dynamics. We again used linear decoders, in this case to predict the past state of the limb, consistent with the anticipated causality with area 2^14,65^ (Methods). As was the case for M1, the low and high frequency bands provided the best predictions (Figure S7B,C). Yet, contrary to M1 (though similar to PMd), the strength of the correlation between an LFP band and the latent dynamics did not predict how well that band could be used to decode movement (Figure S6D). Therefore, we have found robust region-specific LFP-latent dynamics correlation profiles that remain stable for different processes underlying behaviour (Figure 3B, 6B, 7B; Figure S7), and whose correspondence to behavioural decoding is strongest for M1.

### LFP-latent dynamics correlations are not predicted from single neuron features

Our results indicate stable, region-specific, and frequency-dependent relationships between LFP bands and latent dynamics. However, the observed correlations could simply have arisen due to trivial relationships between LFP and single neuron spiking activity^31,40,43,47–50^, rather than being dependent on population-wide latent dynamics. To directly address this, we asked whether LFP bands that were well correlated with the latent dynamics mostly captured the firing rates of the neurons recorded on that electrode. If that were the case, the correlations between LFPs and the firing rate of neurons on the same electrode (“within-electrode correlation”) should exceed the analogous correlations between LFPs and the firing rates of neurons on different electrodes (“across-electrode correlation”) (Methods). In fact, most within-electrode correlations for M1, PMd and area 2 were very low, statistically no different from the across-electrode correlations (Wilcoxon rank sum test, *P* ≥0.1, except for M1 LMP (*P*=0.03) and 0.5–4 Hz (*P*=0.05); Figure 8A), suggesting that the observed similarities between LFPs and latent dynamics cannot be predicted from the single neuron activity.

**Figure 8:**
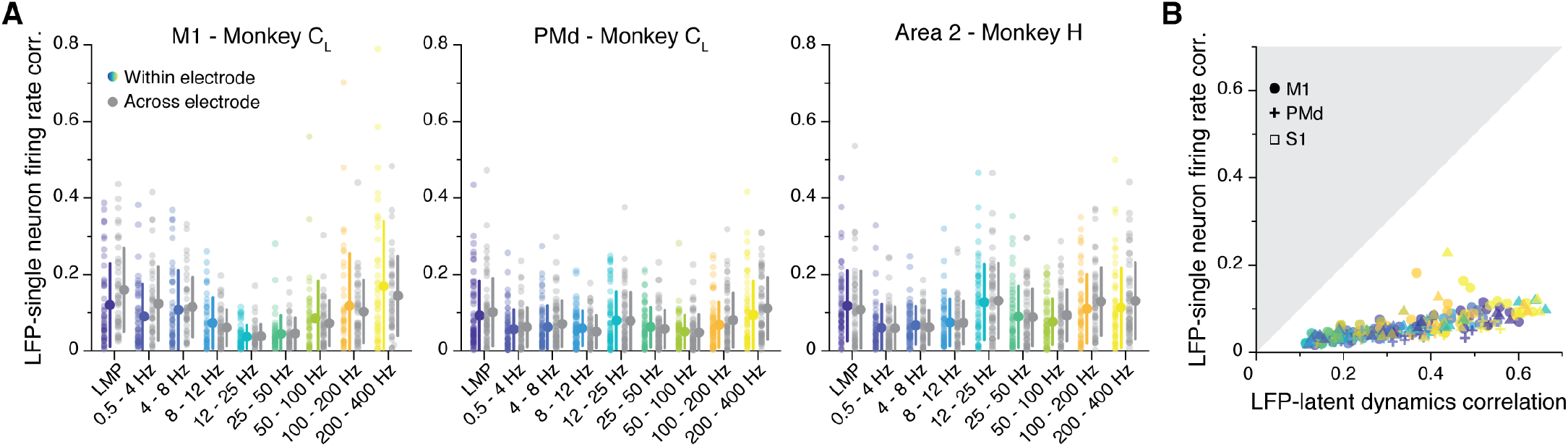
Single neuron features do not account for the LFP-latent dynamics correlations. **A.** Comparison between the correlation of each LFP band with the firing rate of single neurons on the same electrode (coloured), and the activity of single neurons on a different electrode (grey). Each plot shows a representative session from one monkey implanted in each of the three studied regions. Error bars, median ± s.d. **B.** Comparison between the LFP-latent dynamics correlations and the correlation between the LFP and the firing rate of neurons on the same electrode. Data from all sessions and monkeys, shown using region-specific markers (legend). Markers are colour coded according to frequency as in A. Note how the LFP-latent dynamics correlations are much stronger than the correlations between LFPs and the activity of neurons on the same electrode.

As a second control, we compared the LFP-latent dynamics correlation directly to the correlation between the LFP and the single neuron activity on that electrode. Notably, the LFP-single neuron correlations were always much lower than the LFP-latent dynamics correlations (Figure 8B). This was also the case when we “denoised” the single neuron firing rates by subtracting their projection outside the neural manifold, seeking to eliminate the components of the single neuron activity that are not shared across the population (Methods) (Figure S8A-C). Combining all the neurons on the same electrode as “multi-units” also did not change the results (Figure S8D,E). These controls show that specific LFP bands capture population-wide features of the latent dynamics that contribute to their generation in an area-specific, stable manner.

## Discussion

LFPs are intriguing neural signals: changes in LFP power at specific frequency bands correlate well with behavioural and cognitive processes such as initiating a movement^35,40,45,46,80^ or choosing among different stimulus-response rules^28^, yet their relationship with the activity of the neurons driving these processes has long been elusive. LFP biophysics are determined by circuit architecture^7^ and the correlation among synaptic inputs to neural populations^8^. Since both of these also help define the neural covariance structure and shape the latent dynamics, we hypothesised that there should be a fundamental relationship between LFPs and latent dynamics. Here we report, for three different regions of monkey sensorimotor cortex, that associations between LFPs and latent dynamics are stable and frequency-dependent (Figure 3B, 6B, 7B; Figure S7). They are also region-specific, as expected if circuit architecture played a role in determining them.

### Relation to previous single neuron studies

Previous studies investigated the relationship between single neuron activity and LFP power within different frequency bands. Spanning a broad range of regions, from primary visual to motor cortices, they present a puzzling story in which only the high frequency LFPs (typically, 40–80 Hz) are associated with the activity of single neurons or multi-units^31,40,43,47–50^. The lack of single neuron-LFP correlations at low frequencies seems at odds with biophysical models predicting shared synaptic inputs—expected to drive single neuron activity—to be most strongly represented in the low-frequency LFPs^82^. Studies of M1 multielectrode LFP recordings align better with this prediction, since both the high-frequency and low-frequency LFPs allow muscle activity^38^ and movement kinematics^37,39^ to be decoded accurately (Fig. 4). Assuming that a large component of the firing of M1 neural populations relates to the motor commands, the ability to predict behaviour from the low-frequency LFPs hints at a relationship between the two. Such a relationship has been recently modelled as a “mode” that captures behaviour-related dynamics that are shared between the low and high frequency motor cortical LFPs and the neural population activity^83^.

Here, we present a novel approach that explicitly quantifies the relationship between each LFP band and the population-wide latent dynamics. This allows us to show, directly, that low frequency LFP bands are indeed correlated-with the latent dynamics, for each of primary motor (Fig 3B), dorsal premotor (Fig. 6B), and somatosensory cortices (Fig. 7B). We also extend our investigation beyond movement execution by studying motor planning, to demonstrate that the LFP-latent dynamics correlation profiles remain stable throughout different processes of behaviour (Fig. 5A,B, 6C). This result constitutes an important control. Since there is no overt movement during the planning epoch, the robust LFP-latent dynamics correlations during the instructed delay period indicate that their association is not only driven by large common inputs reflecting motor commands: it extends to more cognitive processes such as planning an action. Reassuringly, the LFP-latent dynamic correlations drop during the inter-trial intervals, when monkeys are not engaged in the task and the synaptic currents across the neural population presumably become much weaker, as suggested by the decrease in single neuron firing rates (Figure S4A,B) and the disappearance of large population-wide covariation patterns (Figure S4C). This drop suggests that the LFP-latent dynamics correlations are a consequence of strong, coordinated synaptic currents across the circuit. Interestingly, such a decrease in correlation during the inter-trial period in PMd was not as marked as in M1, perhaps because PMd recapitulates aspects of the task, as shown for PFC^84^. In fact, PMd activity but not M1 activity reflects the probability of possible upcoming reach directions^85^; it also has stronger reward signals than M1^86^.

### A neural population view is necessary

A critical aspect of our results is that the stable LFP-latent dynamics correlation profiles cannot be predicted based on the relationship between LFPs and single neurons. We interpret this based on the importance of circuit architecture in the generation of LFPs^7,9,10^. Biophysical models suggest that strong LFP signals may require correlated synaptic inputs to the neurons generating the electric fields, with the spatial extent of such correlated inputs determining the magnitude of the LFP^8^. These correlated inputs will lead to correlated neural firing patterns that will be captured in the latent dynamics computed with PCA, but much less so in the discharge of any given neuron. Thus, our finding of robust LFP-latent dynamics correlations is likely driven by the biophysical mechanisms of LFP generation and requires explicitly computing the dominant neural covariance patterns. In contrast, the activity of individual neurons is determined by a highdimensional set of inputs affecting any given neuron through unknown weights, blurring their relationship with the LFP.

For a given level of correlated synaptic input, the LFP propagates in a frequency dependent manner, with its spatial reach decreasing quite markedly at higher frequencies^82^. Yet, high-frequency components, especially broadband signals at frequencies >50 Hz, seem also to reflect local neural spiking^48^. Thus, it is possible that the robust associations defining our roughly V-shaped LFP-latent dynamics correlation profiles (Figure 3B, 6B, 7B) have two different sources: at low frequencies, they may be dominated by the correlated synaptic inputs leading to the emergence of low-dimensional latent dynamics in the sensorimotor cortices during behaviour; at high-frequencies, they may reflect a combination of correlated synaptic input across the population and more local neural spiking. In agreement with this, “multi-unit” firing rates at 100-200 Hz and 200-400 Hz are more strongly correlated to the LFPs than single neuron firings rates^37,40^ (compare Figure 8B and Figure S8E), although still less so that the LFP-latent dynamics (Figure S8E).

### Circuit architecture influences the differences between areas

The importance of circuit architecture in determining the LFP properties may also explain the differences in LFP-latent dynamics correlations across areas. Likely due to their varying evolutionary past^87^, each of the three cortical areas we studied has different neuron types, layer organisation and local and long-range connectivity patterns^88^, all of which influence the biophysics of LFP generation^7–9,82^. Because of this, it is reasonable for the “mapping” between latent dynamics and LFPs captured by our correlation profiles to be area dependent.

The most striking difference in LFP-latent dynamics correlation profiles across areas appears at the mid-range 12–25 Hz band. Surprisingly, there is not a clear rostro-caudal gradient in their relationship; instead, the correlation is high in PMd (Figure 6B) and even higher in area 2 (Figure 7B), but extremely low in the intermediately located M1 (Figure 3B). What is common across all three regions is that the 12–25 Hz band is never a good predictor of movement kinematics, even when the LFP-latent dynamics correlations reach 0.5–0.6 (Figure 4C, 6C; Figure S6B,C). This lack of correlation with movement parameters is consistent with studies of M1 LFPs during movement^35,51^, which describe a marked decrease in LFP 12–25 Hz power prior to movement initiation^35,40,45,46^, with some transient oscillations only during specific tasks. These oscillations appear often when monkeys pick up treats, explore their surroundings, or perform relatively challenging finger tasks, but are very rare during more “automatic” wrist tasks^35,41,51^ —the latter being most similar to the centre out reaching task we studied. Combined, these observations suggest that the 12–25 Hz LFP band may be a by-product of population activity related to attention^51^ or other high-level cognitive features such as keeping track of task structure. As such, 12–25 Hz LFP power could potentially be driven by the activity of neural populations in regions projecting to the sensorimotor cortices, rather than by the activity of local populations^89^. In fact, LFP 12–25 Hz power in prefrontal cortex modulates as monkeys select between stimulus-response rules^28^, suggesting that PMd 12–25 Hz oscillations could be correlated with task structure. Analogously, parietal cortical LFP 12–25 Hz power changes as monkeys choose between movement types based on visual cues^32^, again suggesting a potential origin of area 2 12–25 Hz oscillations related to task structure.

### Conclusion

A wealth of studies spanning the entire brain have reported on frequency dependent changes in LFP power that correlate with features of behaviour. Yet, the relationship between the LFPs and the activity of neurons driving behaviour has long remained elusive. Here, we show that, likely due to the biophysics of LFP generation, LFPs are fundamentally related to the shared patterns that dominate the activity of a neural population, rather than to the activity of the neurons themselves. For a given region of the sensorimotor cortex, this association is both stable and frequency dependent.

We anticipate our approach will uncover stable LFP-latent dynamics correlations not only within the sensorimotor cortices but beyond, as low-dimensional latent dynamics have been found across many cortical and subcortical brain regions^12,70^. We further expect to uncover stable LFP-latent dynamic correlations during cognitive tasks, since the two are correlated during “abstract” movement planning. Identifying these relationships offers an exciting opportunity to integrate studies based on either type of signals, including a large body of field recordings in neurological patients^90–97^.

## Acknowledgments

This work was supported in part by: grant F31-NS092356 from the National Institute of Neurological Disorder and Stroke, and grant T32-HD07418 from the National Institute of Child Health and Human Development (M.G.P.); graduate research fellowship DGE-1324585 from the National Science Foundation, and grant T32-NS086749 from the National Institute of Neurological Disorder and Stroke (R.H.C.); grants NS053603, NS074044 and NS095251 from the National Institute of Neurological Disorder and Stroke (L.E.M.); and grant 2017-T2/TIC-5263 from the Community of Madrid, grant PGC2018-095846-A-I00 from the Spanish Ministry of Science and Innovation, and grant EP/T020970/1 from the UKRI Engineering and Physical Sciences Research Council (J.A.G.).

## Methods

### Behavioural task

Four monkeys (males, *macaca mulatta*) were trained to sit in a primate chair and perform a centre-out reaching task using a planar manipulandum. All monkeys completed the task with the hand contralateral to the implanted hemisphere. During the task, the monkey started a trial by bringing the cursor to a target in the centre of the workspace. After a variable waiting period, the monkey was presented with one of eight outer targets (four for Monkey L), which were equally spaced along a circle of 6 to 8 cm radius. Monkeys C and M were trained to wait for the auditory go cue in the centre during a variable delay period of 0.5–1.5 s in which the target remained visible. Monkeys H and L were not subjected to this delay period. To receive a liquid reward, the monkey had to move the cursor into the outer target within 1 s; Monkeys M and C were required to hold the cursor there for 0.5 s, whereas Monkeys H and L were only required to hold for 0.1 s, to ensure they had decelerated and ended the trial within the target. To start a new trial, the monkeys had to return the cursor to the central target. During the task, the endpoint position of the manipulandum was recorded at a sampling frequency of 1kHz; the timing of the task events was digitally logged. Hand velocity was computed as the derivative of the hand position. In all the analyses, we only considered successful trials (an average of 307 ± 221 trials per session; mean ± s.d.).

### Neural implants and recordings

All surgical and experimental procedures were approved by the Institutional Animal Care and Use Committee (IACUC) of Northwestern University. We recorded the spiking activity of neural populations and intracortical LFPs using chronically implanted 96-channel Utah electrode arrays in different regions of cortex. Monkey M was implanted in the M1 and PMd of the right hemisphere. Monkey C was implanted two times: first, he received an array in the right M1 (denoted by C_R_ throughout the text) and then, in a second procedure, he received implants in the left M1 and PMd (denoted by C_L_). Monkeys H and L were implanted in area 2 of primary somatosensory cortex of the left hemisphere.

Neural activity was acquired using a Cerebus system (Blackrock Microsystems, Salt Lake City, UT) at 30 KHz sampling frequency. The recordings on each channel were band-pass filtered (250–5000 Hz), and then converted to spike times based on threshold crossings. The threshold was selected according to the root mean squared (RMS) activity in each channel (Monkeys C and M: 5.5×RMS; Monkeys H and L: 5×RMS). The spike times were later sorted to identify putative neurons using specialized software (Offline Sorter v3, Plexon, Inc, Dallas, TX). The number of identified neurons varied across cortical area (M1: 60 ± 20; PMd: 128 ± 52; S1: 47 ± 25; all mean ± s.d.). We simultaneously recorded the local field potential (LFP) on each channel at a sampling frequency of 1–2 kHz.

### Behavioural epochs

For each trial, we isolated different epochs in the task to identify different “aspects” of the behaviour, namely movement preparation and execution. These windows were adjusted to the behavioural idiosyncrasies of each monkey; however, our results did not change qualitatively within reasonable changes to these windows. For Monkeys C and M, we used a movement execution window starting 120 ms before movement onset and ending 420 ms after onset. For monkeys H and L, the movement execution window started at the go cue, and finished 600 ms after it. For the monkeys subjected to an instructed delay period (Monkeys C and M), we defined a movement preparation window starting 390 ms before movement onset and finishing 60 ms after movement onset. Finally, we also examined neural activity during the inter-trial period –when monkeys did not move the manipulandum– to study the relationship between neural population activity and LFP when monkeys are not engaged in a behaviour using a window starting 540 ms before target presentation and finishing at target presentation.

### LFP processing

The LFP signals were first de-referenced by computing their common average, and then centred at zero by subtracting their average over time. Power line interference was removed using a notch filter (zero-phase Butterworth, 2^nd^ order, f_c_ = 60 Hz). These pre-processed LFPs were then filtered (zero-phase Butterworth, 3^rd^ order) at the following frequency bands: 0.5–4 Hz, 4–8 Hz, 8–12 Hz, 12–25 Hz, 25–50 Hz, 50–100 Hz, 100–200 Hz, and 200–400 Hz. The bandpower of the LFP filtered at each frequency band was computed with an overlapping 50 ms window and 10 ms step. Since a 50 ms window does not cover the whole period of the frequency bands with minimum frequency less than 20 Hz, the band-power at those frequency bands was computed using a window size of 1/f_min_ seconds, maintaining the 10 ms step. All LFP pre-processing procedures follow Ref. 39. We also computed the so-called local motor potential (LMP) by calculating the moving average of the LFP (de-referenced, detrended, and after removing the power line interference) using the same window setting as for the band-power. Note that our results held when we varied the LFP frequency bands within a reasonable range (Figure S2B; details in “Control analyses” below).

Although we recorded LFPs from all 96 array channels in the arrays, we only considered in the analyses the LFPs from channels containing at least one putative neuron clearly identified through manual spike sorting, resulting in 46 ± 16 (mean ± s.d.; ranging from 15 to 77) LFP channels per session. We subsampled the LFP signals to 30 ms bins, to match the bin size used for the single neuron activity, and smoothed them using a Gaussian kernel (s.d.: 50 ms) as we did for the single unit firing rates (see below).

### Neural population latent dynamics

To characterise the dynamics of the neural population activity, we first computed the smoothed firing rate for each putative single neuron by applying a Gaussian kernel (s.d.: 50 ms) to the binned square-root-transformed firings (30 ms bins). Although we excluded neurons with low firing rate (<1 Hz mean firing rate across all bins), we did not perform any additional preselection, e.g., based on directional tuning. For each behavioural epoch^98^ (execution or planning, when present) in each session, this produced a neural data matrix **X** of dimension *n* by *T*, where *n* is the number of neurons and *T* is the total number of time points from all concatenated trials; thus, *T* is the number of trials per session χ the number of points per trial.

We represented the activity of the *n* identified neurons during an epoch in a session in an *n*-dimensional neural state space, in which each axis represents the smoothed firing rate of one neuron. Within this space, we computed the low dimensional manifold that spans the dominant neural covariation patterns^12,72^ by applying Principal Component Analysis (PCA) on **X**. We defined an *m*-dimensional neural manifold by keeping only the leading *m* principal components. Based on previous work by our group and others^14,17,98^, the number of components *m* considered for each area were: *m*=10 for M1, *m*=15 for PMd, and *m*=8 for area 2; for those sessions with an instructed delay period, we used the same manifold dimensionality during the planning and execution epochs. Note that our results held across a wide range of dimensionality values *m* (Figure S1D). We computed the neural population latent dynamics by projecting the smoothed firing rates of the *n* neurons onto the neural manifold. This produced a data matrix **L** of dimension *m* by *T*, where *m* is the dimensionality of the manifold.

### Alignment of latent dynamics and LFPs

We hypothesised that there should be an association between LFPs and latent dynamics, and that such an association would be region-specific, frequency-dependent, and stable during movement planning and execution. To address these hypotheses, we computed the similarity between the latent dynamics and the time-varying changes in LFP power in each band for each behavioural epoch, monkey, session, and cortical area. We used Canonical Correlation Analysis (CCA), a method that quantifies the similarity between two sets of multidimensional signals by finding the linear combinations that, applied to each set, maximally correlates the resulting projections^74^; we refer to this process as “aligning” an LFP band and the latent dynamics.

We start the alignment procedure by considering the concatenated latent dynamics, **L**, and the concatenated LFP power in one electrode at one frequency band *b*, **F**_*b,i*_, where *b* is one of the nine frequency bands listed above (including the LMP), and *i* the electrode number; the vector **F**_*b,i*_ thus has dimension 1 by *T*. For each LFP band, we align separately the latent dynamics and the LFP power at one frequency band in each of the *k* electrodes with at least one putative neuron, which yields *k* canonical correlation coefficients (CCs) per LFP band. Throughout the paper, we summarise these distributions based on their mean ± s.d. (see, e.g., Figure 3B, 6B, 7B). For details on the implementation of CCA, see our previous work, Ref. 14.

To test whether the CCs between each LFP band and the latent dynamics capture a significant association between these two types of signals, we devised a control using Tensor Maximum Entropy (TME), a method that generates surrogate neural population data preserving selected features of the original data^76^. Here, we used TME to generate surrogate neural firing rates that preserved the covariance across neurons but that had different covariance across trials and over time (i.e., different dynamics) than the actual data. We then smoothed these surrogate firing rates with a Gaussian kernel (s.d.: 50 ms), and applied PCA to obtain a matrix of surrogate latent dynamics, which were very similar to the actual latent dynamics based on their spectral content (Figure S1B,C). To assess whether the experimentally measured CCs captured a significant association between an LFP band and the latent dynamics, we compared them to the CCs obtained after aligning the same LFP signals to the surrogate latent dynamics (shown in colour and in black, respectively, in Figure 3B, 6B, 7B).

### Decoding hand velocity from M1 and S1 movement activity

After quantifying to what extent each M1 and S1 LFP band correlates with the latent dynamics, we asked how well it predicts behaviour. We trained Wiener filters^99^ to predict hand velocity from each of the nine LFP bands, including the LMP. The filters included three bins of recent LFP history, for a total of 90 ms. These additional inputs seek to account for the intrinsic LFP dynamics and transmission delays. Given that M1 activity causes movement, for this region we included additional bins preceding the hand velocity signals. In contrast, since S1 activity is largely driven by responses to the ongoing movement, for this region we included additional bins lagging the hand velocity signals. We further added a 3^rd^ order polynomial at the output of the filter to compensate for nonlinearities^100^. Finally, to obtain accuracy values against which to compare the LFP decoders, we computed similar Wiener filters using the M1 or S1 latent dynamics as inputs. Decoder prediction quality was measured based on the *R*^2^ between the actual and predicted velocities. We built separate decoders for the X and Y hand velocities; as they had similar accuracy, we report their mean.

To avoid overfitting, we used 90% of the trials in the session to train the decoder and the remaining 10% to test it. We repeated this procedure 50 times, randomly selecting the non-overlapping training and testing trials in each iteration. We averaged the *R*^2^ value for the X test blocks to obtain the final reported value.

### Predicting reach direction from PMd planning activity

We trained naïve Bayes classifiers^63^ to predict the direction of the upcoming reach based on the pre-movement LFP activity in PMd. Similar to the velocity decoders, we built nine sets of LFP-based classifiers, each taking the LFP signals at one frequency band as inputs, as well as latent dynamics-based classifiers whose performance we used as reference. In all cases, we used PMd activity within a 450-ms window as input to the classifier. Within this window, we averaged all LFP activity (or latent dynamics) to obtain a single bin per trial, and then standardized it using a z-transform. Prediction accuracy was measured as the percentage of targets predicted correctly.

To protect against overfitting, we trained the classifier in 90% of the trials and tested the performance on the remaining 10%. As for velocity decoders, we repeated the procedure 50 times randomly selecting the non-overlapping training and testing trials in each iteration. We averaged the percentage of correct predictions to obtain the final reported value.

### Control analyses

We performed all the analyses in the paper using the standard LFP bands describe above. In addition, we also verified that our alignment results held even if the LFPs were filtered in different bands by repeating the analysis on partly overlapping bands that swept the entire LFP spectrum (Figure S2). We chose pre-processing frequency bands that had width similar to that of the “classic” LFP bands, which resulted in 53 bands with increasing width, ranging from 4 to 100 Hz. Filtering windows had a 50 % overlap, and the following widths: 4 Hz between 0.5 and 14.5 Hz, 12 Hz between 10.5 and 26 Hz, 25 Hz between 12.5 and 57.5 Hz, 50 Hz between 25 and 120 Hz, and 100 Hz between 50 and 440 Hz.

We devised two control analyses to test whether the similarities between LFP bands and latent dynamics could be explained by features of the single neuron activity. In the first control, we verified that the LFP on an electrode does not strongly reflect the activity of single neurons in that electrode. Thus, we verified that the relationship between LFP bands and single neurons on the same electrode does not exceed that between LFP bands and single neurons on different electrodes. We computed the Pearson correlation between the activity of each putative neuron and the power of each LFP band on that same electrode; this yielded nine distributions of *n* correlation values. We then computed the Pearson correlation between each LFP band and the activity of a randomly selected neuron on a different electrode; this again yielded nine distributions of *n* correlation values. Note that neurons detected on neighbouring electrodes were excluded to make sure the distance in the across-electrode group was not small. Direct comparisons between these paired sets of distributions allowed us to show that single neuron activity does not explain away our results (Figure 8A). As a second control, we compared directly, for each LFP band, the correlation between the LFP and the single neuron activity to the CC between the LFP and the latent dynamics (Figure 8B).

We repeated these controls after summing up the spikes of all putative single neurons on an electrode to form a “multi-unit” (Figure S8D,E), and after “denoising” the single neuron firing rates based on PCA (Figure S8A-C). This denoising consisted in subtracting from the smoothed firing rate of each neuron its projection outside the neural manifold, i.e., the projections onto dimensions *m*+1 to *n*. Notably, none of these manipulations changed the results qualitatively.

## Data availability

The datasets analysed for this manuscript will be shared upon reasonable request.

## Code availability

All analyses were implemented using custom Matlab (The Mathworks Inc.) code. Code to replicate the main results will be shared upon reasonable request.

## Supplementary Figures

**Figure S1:**
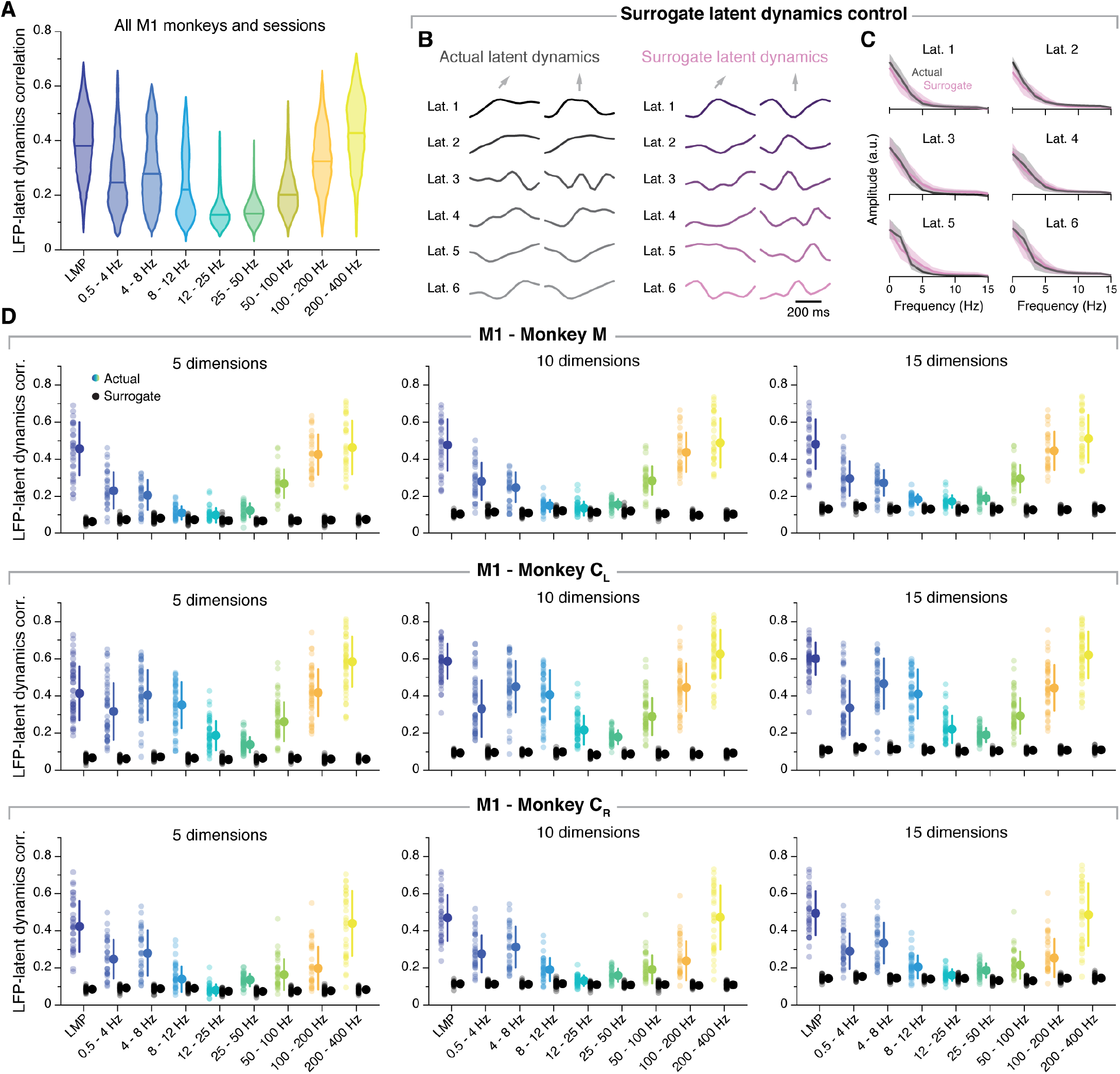
Additional data: Similarity between M1 latent dynamics and each LFP band, and control analyses. **A.** Correlation between each LFP band and the latent dynamics; data pooled across all M1 sessions and monkeys. Note that the correlation profile matches well the example sessions in Figure 3. Violin, probability density for one frequency band; horizonal line, median. **B.** Example actual latent dynamics and surrogate latent dynamics generated using TME for two randomly selected trials from the example session from monkey C_L_ shown in Figure 3. Only six of ten dimensions are shown for visualization purposes. **C.** Amplitude spectrum of the actual and surrogate latent dynamics for the same session. Line and shaded areas: mean ± s.d across all the trials in the session. Note the strong similarity in spectral content between actual and surrogate latent dynamics. **D.** Similarity between M1 latent dynamics and each LFP band for three different manifold dimensionalities. The plots show the same representative session from each M1 monkey shown in Figure 3 using the same format. Note that the trend in the LFP-latent dynamics correlation profile does not depend on the assumed dimensionality of the neural manifold.

**Figure S2:**
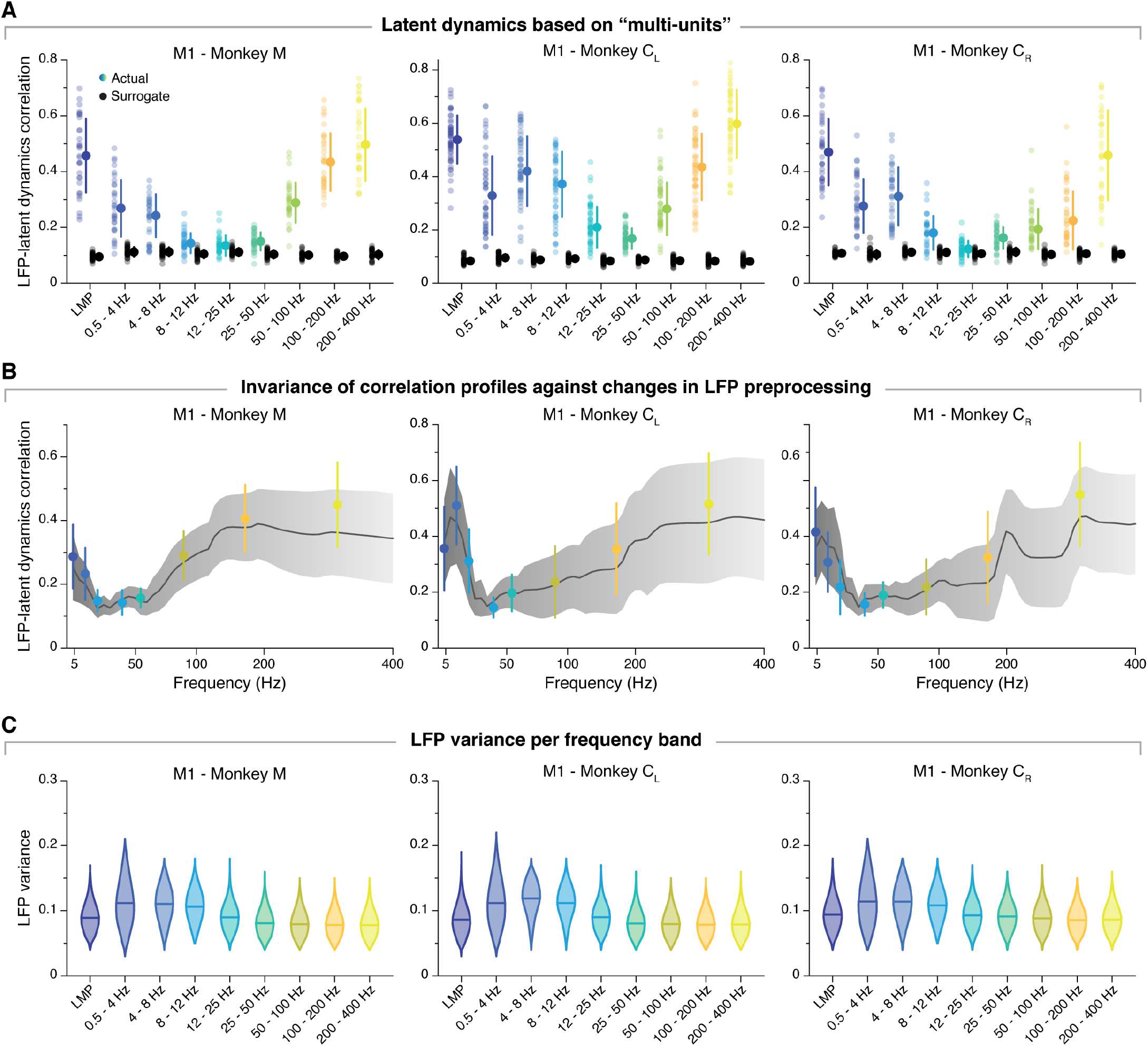
Additional data: Further control analyses to illustrate the similarity between M1 latent dynamics and each LFP band. **A.** Correlation between each LFP band and the latent dynamics (coloured markers) after pooling all single neurons on each electrode as a “multi-unit”. Data for the same representative sessions from each M1 monkey shown in Figure 3 and S1. Black markers show the control correlation values obtained after generating surrogate neural activity using TME. Note that pooling single neurons into “multi-units” does not alter the results: all the trends are the same as in Figure 3B. Error bars, median ± s.d. **B.** The latent dynamics-LFP correlation profile does not depend critically on the frequency bands used for LFP pre-processing; it follows the same trend if the LFP power is computed on partly overlapping bands that sweep the entire signal spectrum (see Methods). The example sessions from each M1 monkey are the same as in Figure 3B (the coloured error bars are reproduced from that same figure). Line and shaded areas, mean ± s.d. **C.** LFP power variance in each frequency band during the same example session from each M1 monkey shown throughout the paper. LFP variance is quite similar across bands, and that the bands with highest correlations with the latent dynamics (LMP, 100-200 Hz, 200-400 Hz) are among those with lowest variance. Violin, probability density for one frequency band; horizonal line, median.

**Figure S3:**
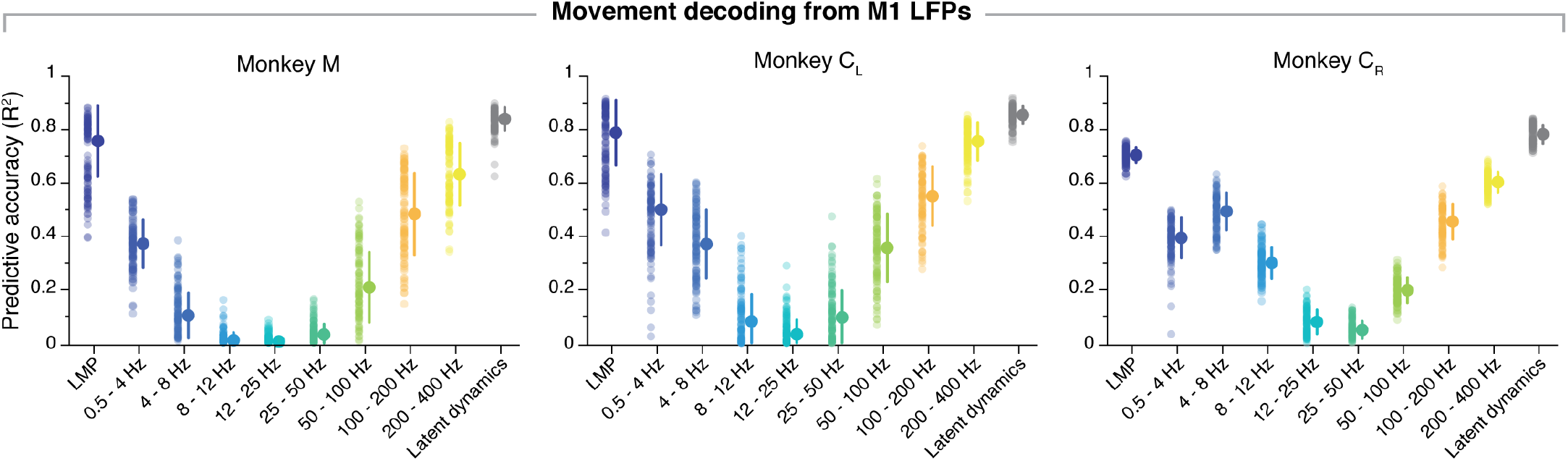
Additional data: Decoding movement kinematics from M1 LFPs. Predictive accuracy of each LFP band (colours) and the latent dynamics (grey) for one representative session from each M1 monkey (sessions are the same used in Figure 3, S1, and S2). Error bars, median ± s.d.

**Figure S4:**
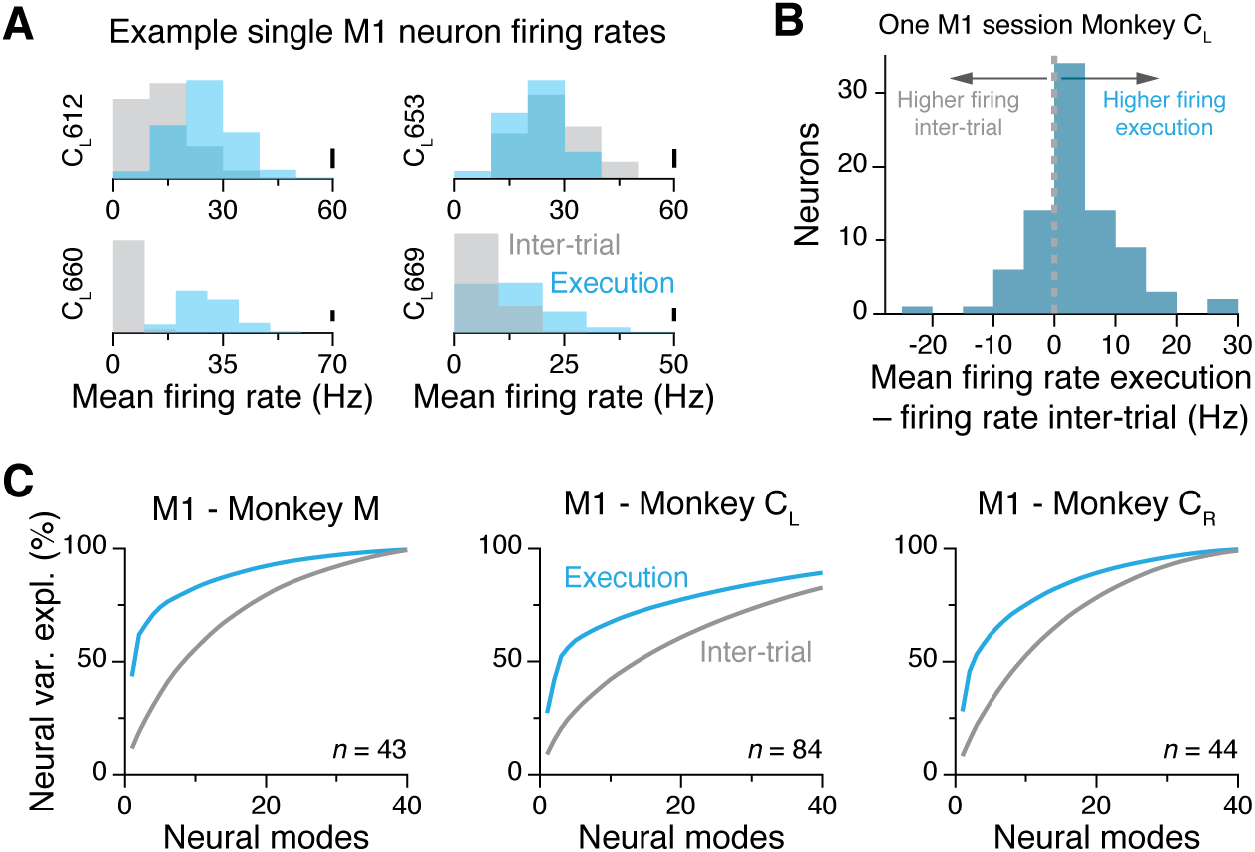
Additional data: Decrease in M1 firing rates and neural correlations when monkeys are not engaged in the task. **A.** Change in mean firing rate between the movement epoch and the inter-trial epoch –when monkeys were not engaged in the task– for four example neurons. Data from the example session from Monkey C_L_ shown throughout the paper. Each data point represents one trial. Scale bar: 20 trials. **B.** Difference in average single neuron firing rates between the movement execution and inter-trial epochs for all 84 single neurons recorded during the same session as in A. As expected, the firing rate of M1 neurons tends to increase when animals are moving. **C.** Percentage neural variance explained by manifolds with increased dimensionality (a “neural mode” is an eigenvector defining the PCA manifold; see Ref. 12). During the inter-trial period, neural manifolds with a given dimensionality *m* explain a considerably lower percentage of total variance than manifolds with the same dimensionality *m* do during movement.

**Figure S5:**
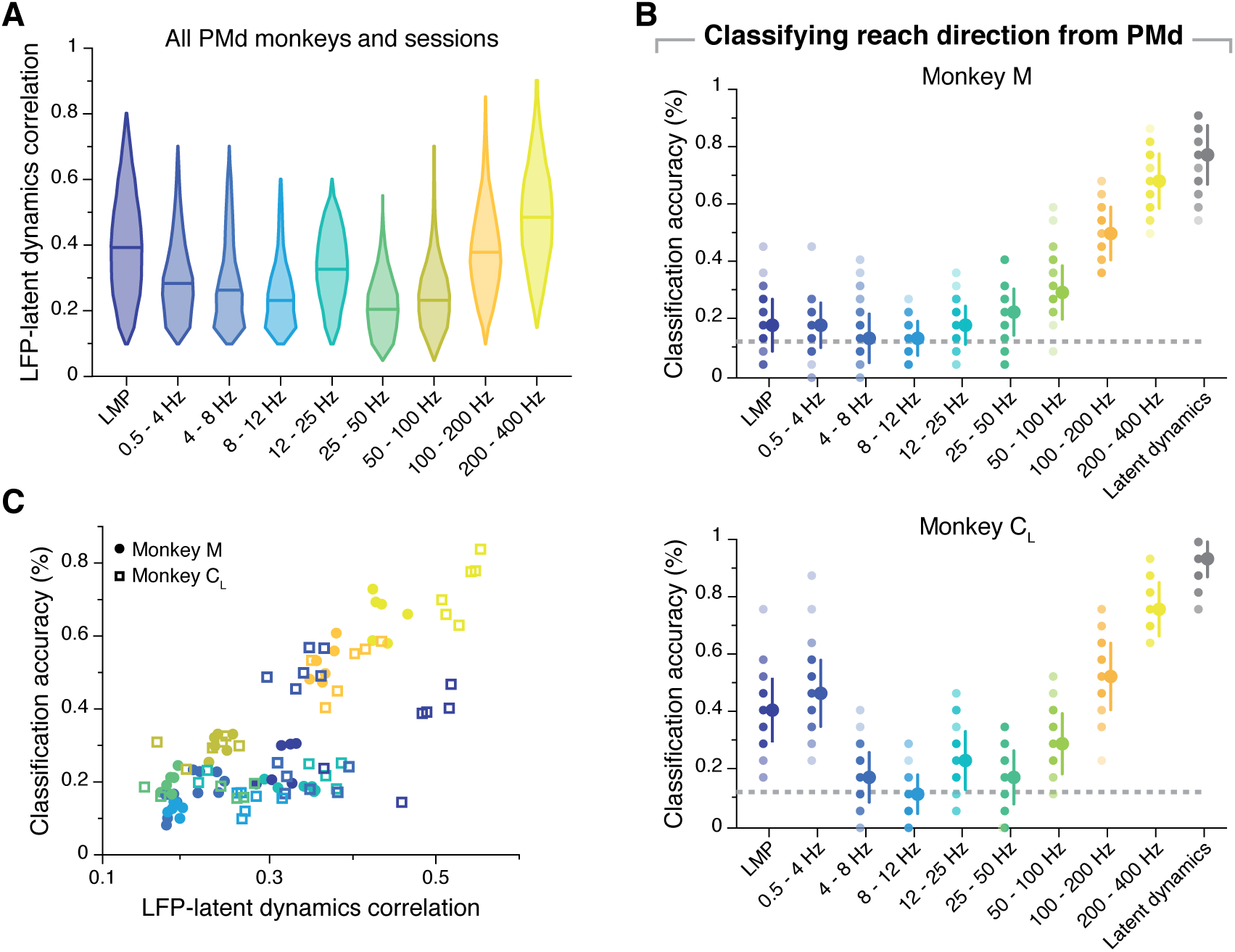
Additional data: Similarity between PMd latent dynamics and each LFP band, and predicting reach direction from PMd LFPs. **A.** Correlation between each LFP band and the latent dynamics; data pooled across all PMd sessions and monkeys. Note that the correlation profile matches well the example sessions in Figure 6. Violin plot, probability density for one frequency band; horizonal line, median. **B.** Accuracy of classifiers that predict the reach direction based on each LFP band (colours) and the latent dynamics (grey) during one representative session from each PMd monkey (same sessions shown in Figure 6). **D.** The relationship between the PMd LFP-latent dynamics correlation coefficients and LFP classification accuracy is not linear, in contrast to M1. Each marker denotes one LFP band from one session; different monkeys are shown using different markers (legend). Markers are colour-coded as in B.

**Figure S6:**
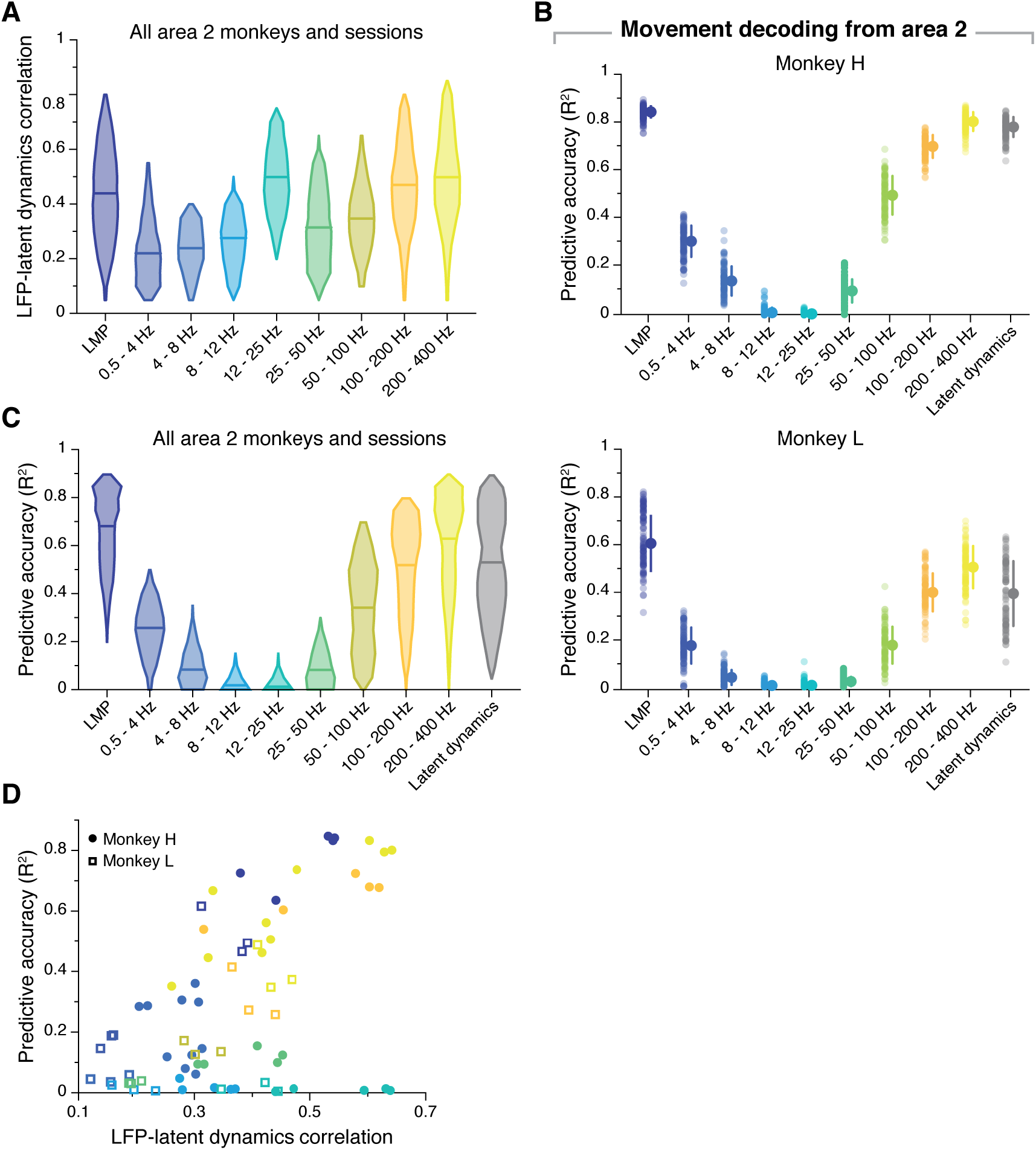
Additional data: Similarity between area 2 latent dynamics and each LFP band, and decoding hand kinematics from area 2 LFPs. **A.** Correlation between each LFP band and the latent dynamics; data pooled across all area 2 sessions and monkeys. Note that the correlation profile matches well the example sessions in Figure 7. Violin plot: probability density for one frequency band; horizonal line, median. **B.** Predictive accuracy of each LFP band (colours) and the latent dynamics (grey) for one representative session from each area 2 monkey (sessions are the same used in Figure 7). Error bars, median ± s.d. **C.** Predictive accuracy of each LFP band (colours) and the latent dynamics (grey) pooled across all area 2 monkeys and sessions. Same format as A **D.** The relationship between the area 2 LFP-latent dynamics correlation coefficients and LFP predictive accuracy is not linear, in contrast to M1, but similar to PMd. Each marker (colour-code as in A) denotes one LFP band from one session; different monkeys are shown using different markers (legend).

**Figure S7:**
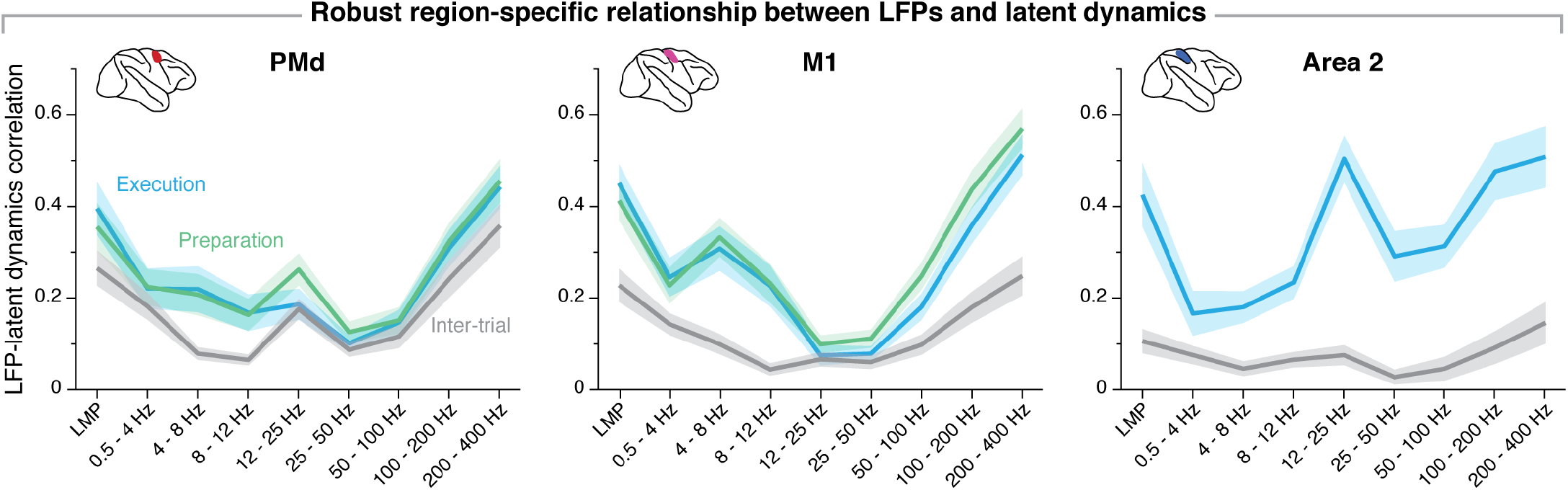
Region-specificity of the LFP-latent dynamics correlation profile. Comparison of the correlation between each LFP band and the latent dynamics across three sensorimotor cortical regions. For both PMd and M1, the correlation profiles remain virtually identical between movement preparation and movement execution. For all three regions, the LFP-latent dynamics correlation profiles decrease drastically when monkeys are not engaged in the task. Line and shaded areas, mean ± s.d. across all sessions from all monkeys for each area. The M1 panel is reproduced from Figure 5A.

**Figure S8:**
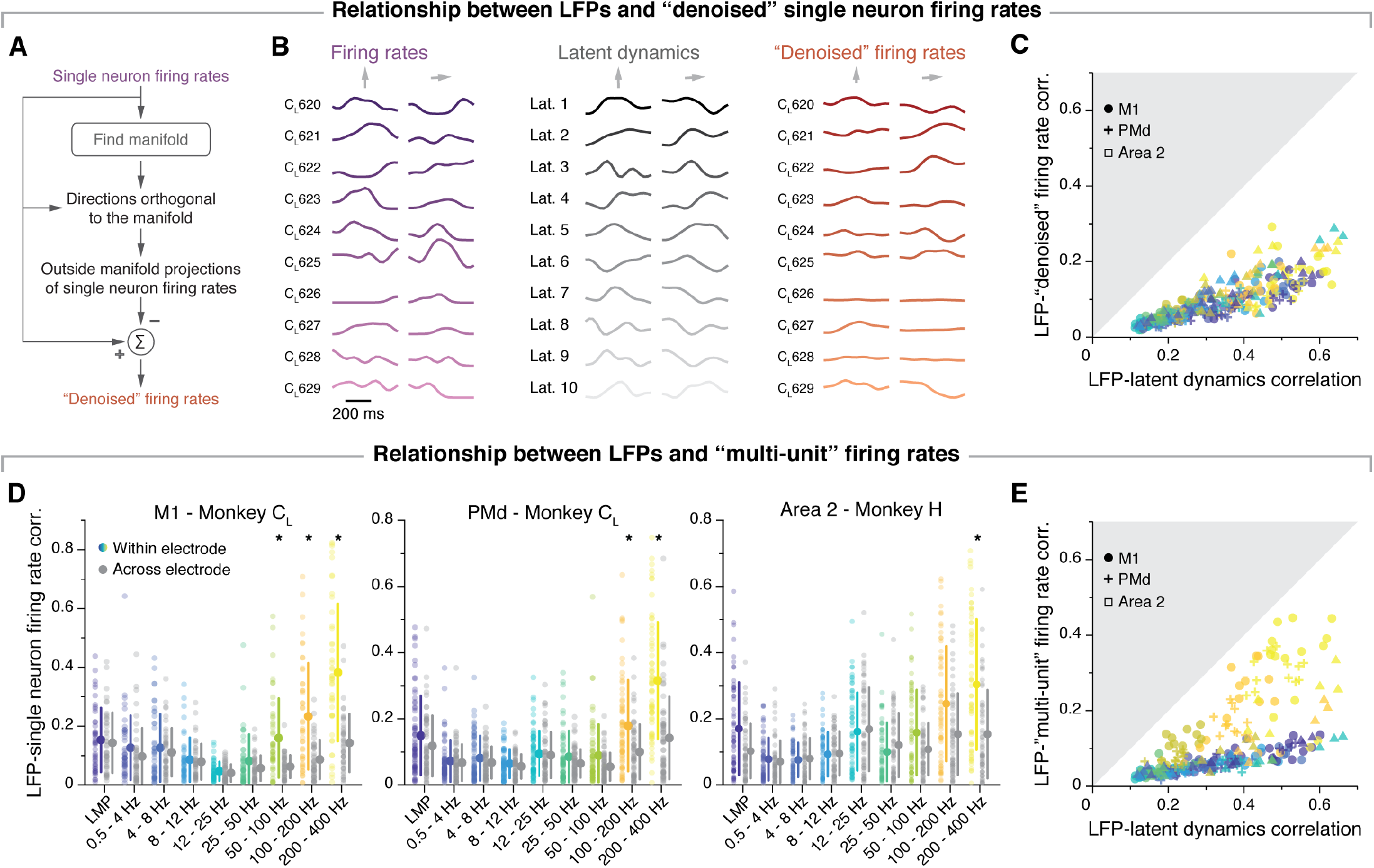
Additional data: Single neuron features do not account for the LFP-latent dynamics correlations. **A.** To match the smoothness of the single neuron firing rates to that of the latent dynamics, we “denoised” the former by subtracting their projections outside the neural manifold (see Methods). **B.** Examples of firing rate “denoising” for two randomly selected trials from one example session from Monkey C_L_. Each of the three columns shows, from left to right: the single neuron firing rates for ten randomly selected neurons, the neural population latent dynamics along each of the 10 manifold dimensions, and the denoised firing rates from the same neurons shown on the left. **C.** Comparison between the LFP-latent dynamics correlations and the correlation between the LFP and the “denoised” firing rate of neurons on the same electrode. Data formatted as in Figure 8B. **D.** Comparison between the correlation of each LFP band with the “multi-unit” activity on the same electrode (coloured), and the “multiunit” activity on a different electrode (grey). As in previous analysis, “multi-units” were obtained by pooled together all single neurons detected on each electrode. Each plot shows a representative session from one monkey implanted in each of the three studied regions. Error bars, median ± s.d. * *P*<0.001 two-sided Wilcoxon’s rank sum test. **E.** Comparison between the LFP-latent dynamics correlations and the correlation between the LFP and the “multi-unit” activity on the same electrode. Data formatted as in C. Note that neither denoising the firing rates, not pooling the activity of single neurons into multi-units affects the conclusion that the neural population latent dynamics are more strongly related to the LFPs than single neurons (or “multi-units”) are.

